# Tomographic docking suggests the mechanism of auxin receptor TIR1 selectivity

**DOI:** 10.1101/081794

**Authors:** Veselina V. Uzunova, Mussa Quareshy, Charo I. del Genio, Richard M. Napier

**Affiliations:** School of Life Sciences, University of Warwick, Gibbet Hill Road, Coventry CV4 7AL, UK

**Keywords:** tomographic docking, auxin, receptor selectivity, molecular filter

## Abstract

We study the binding of plant hormone IAA on its receptor TIR1 introducing a novel computational method that we call *tomographic docking* and that accounts for interactions occurring along the depth of the binding pocket. Our results suggest that selectivity is related to constraints that potential ligands encounter on their way from the surface of the protein to their final position at the pocket bottom. Tomographic docking helps develop specific hypotheses about ligand binding, distinguishing binders from non-binders, and suggests that binding is a three-step mechanism, consisting of engagement with a niche in the back wall of the pocket, interaction with a molecular filter which allows or precludes further descent of ligands, and binding on the pocket base. Only molecules that are able to descend the pocket and bind at its base allow the co-receptor IAA7 to bind on the complex, thus behaving as active auxins. Analyzing the interactions at different depths, our new method helps in identifying critical residues that constitute preferred future study targets and in the quest for safe and effective herbicides. Also, it has the potential to extend the utility of docking from ligand searches to the study of processes contributing to selectivity.

## I. INTRODUCTION

The molecular recognition of specific small organic compounds by target proteins is of central importance in biology. Auxins are a particularly relevant class of small molecule plant hormones with considerable importance for growth and development. Both the naturally occurring indole-3-acetic acid (IAA) and synthetic auxins bind to the Transport Inhibitor Response 1 (TIR1) family of receptors. In turn, auxin binding to the receptor allows the co-receptor IAA7 to bind to the substrate-receptor complex [1, 2]. Thus, one can say that auxins act as “molecular glues” between partners of the receptor system. The completion of this two-step mechanism triggers a cascade of events leading to changes in gene expression [3]. The macroscopic results of acute exposure to synthetic auxins are explosive, epinastic growth followed by plant death. Thus, such compounds have found widespread application in herbicidal formulations. Further valuable features of auxin-based herbicides are a long history of safe use and their selective action against broad-leaved plants, making them preferred products for the control of weeds in cereal crops and turf [4]. However, rational design of novel biologically active molecules to influence the TIR1 receptor has proved challenging because the protein recognizes a diverse set of natural and synthetic ligands [2]. At the same time, TIR1 is highly selective. For instance, the native ligand IAA shares many structural features with its biosynthetic precursor, the indolic amino acid L-tryptophan (Trp), which, although ubiquitous and present at intracellular concentrations far in excess of that of IAA, does not elicit auxin responses [5].

A likely reason for this is that inactive compounds do not bind the receptor in the right location or with the right orientation, if at all, thus precluding assembly of the co-receptor complex. Then, knowledge of the mechanism of interaction for natural ligands is fundamental for the design of synthetic analogues. Computational methods for molecular docking have become standard tools in active compound design and discovery [6–8]. They allow a reduction of the search space, leading to targeted experimental binding assays, and they are widely used for ligand screening and identification of binding sites on bioactive targets [9, 10]. Specific examples of the application of molecular docking are the identification of a genetic cause of cancer drug resistance [11], ligand differentiation between human oestrogen receptors [12], rational drug design for neurodegenerative diseases [13], modelling candidate therapeutic binding to mutated targets [14], and design of highly catalytic artificial metallobioenzymes [15]. In all cases the binding site is considered as a single, holistic search space. In this article, we introduce a new approach to molecular docking, which we call *tomographic docking*, that we use to propose an explanation for the discrimination mechanism of small ligands by the TIR1 receptor.

One frequently overlooked aspect of the molecular recognition process is the depth of the protein binding site, which can extend significantly towards the protein core. Several computational tools exist that help describe and define pockets, tunnels, channels and pores, and some will identify the most likely high-affinity sites in the recess. Once defined, these are offered as binding sites for docking. While this approach is able to find a good candidate for the lowest-energy configuration of a given receptor-binder pair, it risks neglecting receptor features that will be encountered by the ligand on the approach to the best site. When this happens, it contributes to ligand misidentification and false positive results, both of which are recognized issues with docking experiments [16]. For example, AutoDock Vina [17, 18], which is currently one of the most popular docking platforms due to its speed, reliability and output accessibility, finds an apparently viable docking pose for Trp on TIR1, even though Trp is experimentally proven to be a non-binder. It is thus reasonable to consider the passage of a ligand into a deep binding site as a multi-step process composed of many interactions, sequential in time and space. Consequently, typical docking approaches, which consider any geometrically valid pose as equally viable, may overlook important physical and chemical barriers that could preclude some potential binders from accessing an otherwise ideal site. To take into account the entire structure of the deep pocket of TIR1, we create a new docking approach. Rather than considering the whole TIR1 pocket as a single, whole entity, our method divides it into a number of “slices” across its depth. Each slice is treated individually, so that the results we obtain in terms of scoring functions and orientations change progressively with depth. In analogy to the tomographic scans routinely employed for medical diagnoses, we call our method *tomographic docking*. The analysis of a whole series of results allows us to identify physical constraints that preclude the binding of Trp while allowing that of IAA. Also, we identify the structural features of the pocket likely to be responsible for the mechanism of selectivity.

## II. METHODS

### A. The target protein

The X-ray crystal structure of TIR1 is solved in three different binding states: unbound/empty (2P1M), in complex with the natural ligand IAA (2P1P, Fig.1), and assembled with both IAA and its co-receptor IAA7 (2P1Q). For our study, we use the unbound structure 2P1M as the closest approximation to what a free ligand interacts with. Superposition of 2P1M with 2P1P and 2P1Q suggests no significant conformational change is induced by ligand binding [1].

**Figure 1:**
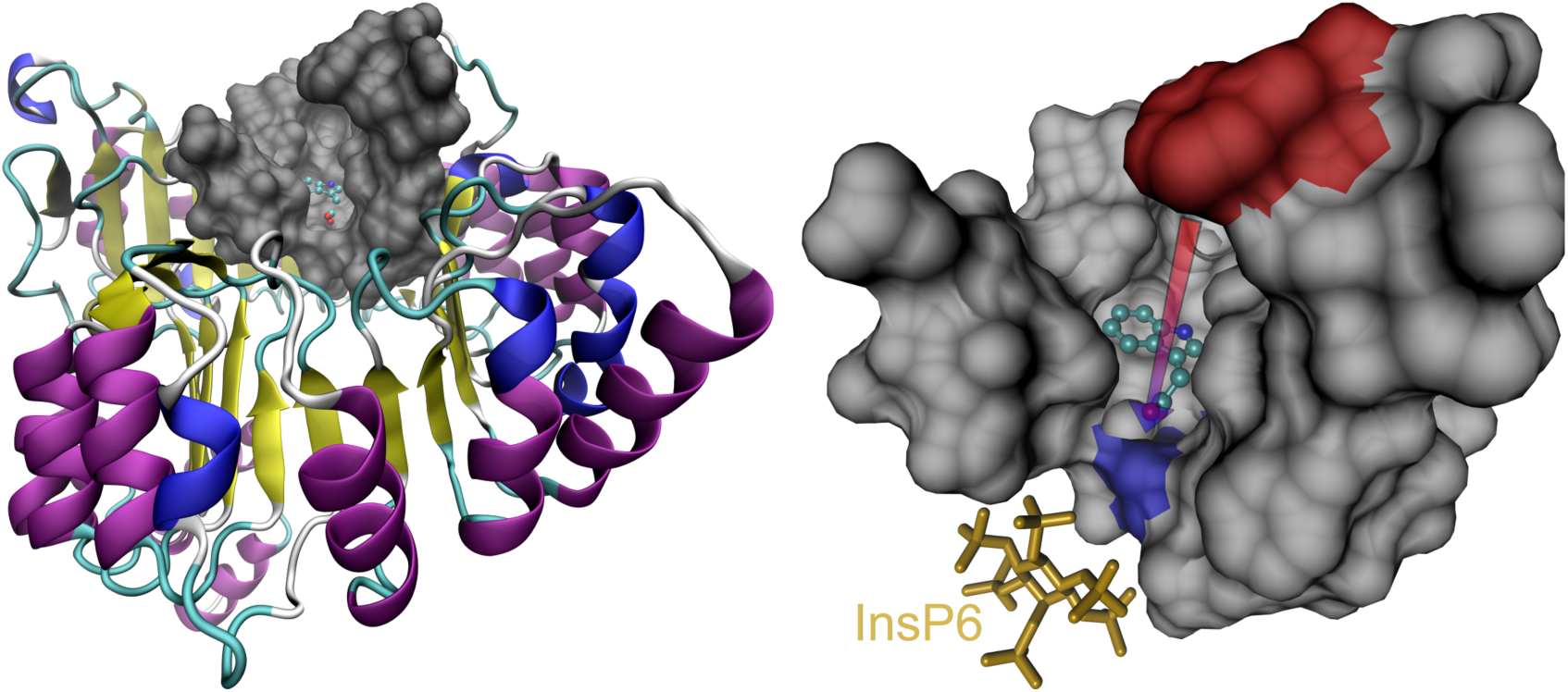
The deep binding pocket on TIR1. The binding site of TIR1 is not a shallow surface indentation but a deep pocket, shown in SURF representation in the left panel superimposed on the cartoon of the whole receptor (2P1P). The residues comprising the binding pocket, isolated and shown from a closer point in the right panel, are contributed by seven, non-sequential leucine-rich repeats [1]. The two reference residues Phe-351, in red, and Arg-403, in blue, indicate the mouth and the bottom of the pocket, respectively. Their distance, indicated by the arrow in the figure and corresponding to the pocket depth, is 16.5Å. IAA is shown in CPK representation, and InsP6 is on the bottom left in bond representation. All 3d molecular visualizations were produced using VMD [21, 22].

The crystallography data contain several associated biomolecules,as well as water. Thus,to prepare the docking input, we first processed the data using VMD [21, 22], excising water molecules and the co-expressed SCF^TIR1^ adaptor ASK1. AutoDockTools [23, 24] was then used to produce the final pdbqt input files. Note that TIR1 harbours inositol hexaphosphate (InsP6) as a second, probably structural, ligand. We left this in place because, unlike ASK1, its location is physically close to the bottom of the pocket. Nonetheless, our final results show that its position is still too far from the binding site to generate effective interactions with the ligand.

### B. Ligands

The investigation focusses on IAA and Trp as TIR1 ligands because Trp, which is a precursor in the synthesis of IAA [25], has no auxin activity and no TIR1- binding activity [5], notwithstanding a significant structural similarity with IAA (Fig. S1). To validate the accuracy of our approach, we later extend the analysis to a few other compounds, comprising both binders and non-binders, with different degrees of structural similarity to IAA (Fig. S2). To prepare the ligands for docking, we first calculated their protonation state at pH 7.3 in water, since binding to the receptor occurs at physiological pH in the plant cell nucleus. Then, we computed their equilibrium geometry using density functional theory with EDF2 functional [26] and a 6-31G* split-valence basis. Finally, we generated the pdbqt files using AutoDock-Tools [23, 24].

### C. Numerical setup

To define the search space to be used in the docking algorithm, we observed SURF representations of TIR1 in both bound and unbound states, and noted that the binding site is not a shallow surface feature, but rather a deep binding pocket (see Fig. 1). In particular, we identified the constituents of the pocket to be a total of 43 amino acids in seven contiguous sets on the leucine-rich repeat loops, namely residues 77–84, 344–354, 377–381, 403–410, 436–441, 462–465 and 489–490. The pocket thus defined has a depth of 16.5 Å, as measured between Phe-351 at the mouth and Arg-403 at the bottom. To investigate the engagement process as the ligand moves into it towards the final binding site at the bottom, we defined an 18 Å× 18 Å× 18 Å cubic search space that moves from above the pocket mouth to below the bottom in steps of 1 Å. The search space at the first step includes Phe-351 at its bottom, and its motion is parallel to the principal axis of inertia of the receptor, whose direction is along the pocket depth. At the last step, Arg-403 is completely included. Then, we performed independent numerical docking experiments at each step, building a sequence of results that provide information on the descent of the ligands into the deep pocket. For the actual simulations, we created a code that automates the tomographic scanning process by computing the geometry of the search space for any specified number of steps, search exhaustiveness, and set of ligands. The code, which we refer to as TomoDock, uses AutoDock Vina [17, 18] as docking engine, and produces tunable summaries of the results, as well as pdb files for further analysis and visualization. Note that with the choices detailed above, the search space is always larger than Vina’s cutoff threshold for the interactions, which is 8 Å. The standard TomoDock experiment was repeated 100 times, with search exhaustiveness of 16.

### D. Experimental setup

Experimental evaluation of the numerical results was carried out using surface plasmon resonance (SPR) as a test of ligand binding to TIR1, and root growth assays for overall biological activity. We performed protein purification and set up the SPR experiments according to the protocols described in [5]. TIR1 was expressed in insect cell culture using a recombinant baculovirus. The construct contained sequences for three affinity tags, namely 6×His, maltose-binding protein (MBP) and FLAG. Initial purification using the His tag was followed by cleavage of His-MBP using TEV protease. After TEV removal and clean-up using FLAG chromatography, the purified protein was used for SPR assays by passing it over a streptavidin chip loaded with biotinylated IAA7 degron peptide. The SPR buffer was Hepes-buffered saline with s1 mM EDTA, 0.05% P20 and 1mM TCEP. Compounds to be tested were premixed with the protein to a final 50 *µ*M concentration. Binding experiments were run at a flow rate of 20 *µ*l/min using 3 minutes of injection time and 2.5 minutes of dissociation time. Data from a control channel (biocytin) and from a buffer-only run were subtracted from each sensorgram following the standard double reference subtraction protocol. To assay root growth inhibition, Col-0 WT seeds were stratified at 4 ◦C for 48 hours on plates containing ½ Murashige and Skoog medium, followed by incubation for 6 days in 12-hour day/night cycles, at a temperature of 20 ◦C for the day and 18 ◦C for the night. Seedlings were then transferred to fresh plates containing test compound and poured fresh on the day. After a further 6 days of growth plates were scanned and the extension of the primary root during treatment was measured using ImageJ [19, 20].

## III. RESULTS AND DISCUSSION

### A. Conventional docking

We first ran a conventional, “static” docking experiment using AutoDock Vina using a cubic search space with an 18-Å side encompassing the whole pocket area. The results show that IAA docks at the pocket bottom and, even from a single run, the best docked position closely matches that of the ligand bound in the co-crystallised structure (Fig. 2). The indole ring is aligned parallel to the pocket bottom and is nested in a semicircle of four non-polar residues, while the carboxylate anion orients itself towards a group of basic residues with which hydrogen bonds are made [1]. Despite the absence of activity for Trp as an auxin and no measurable affinity for binding, AutoDock Vina finds an apparently plausible docked position for it at the bottom of the pocket, although not with the same orientation as IAA (Fig. 2). This clearly shows that one cannot rely on a direct interpretation of docking results to identify binders, because a “cover-all” search space overlooks key features of the binding process and a more systemic approach is needed.

**Figure 2:**
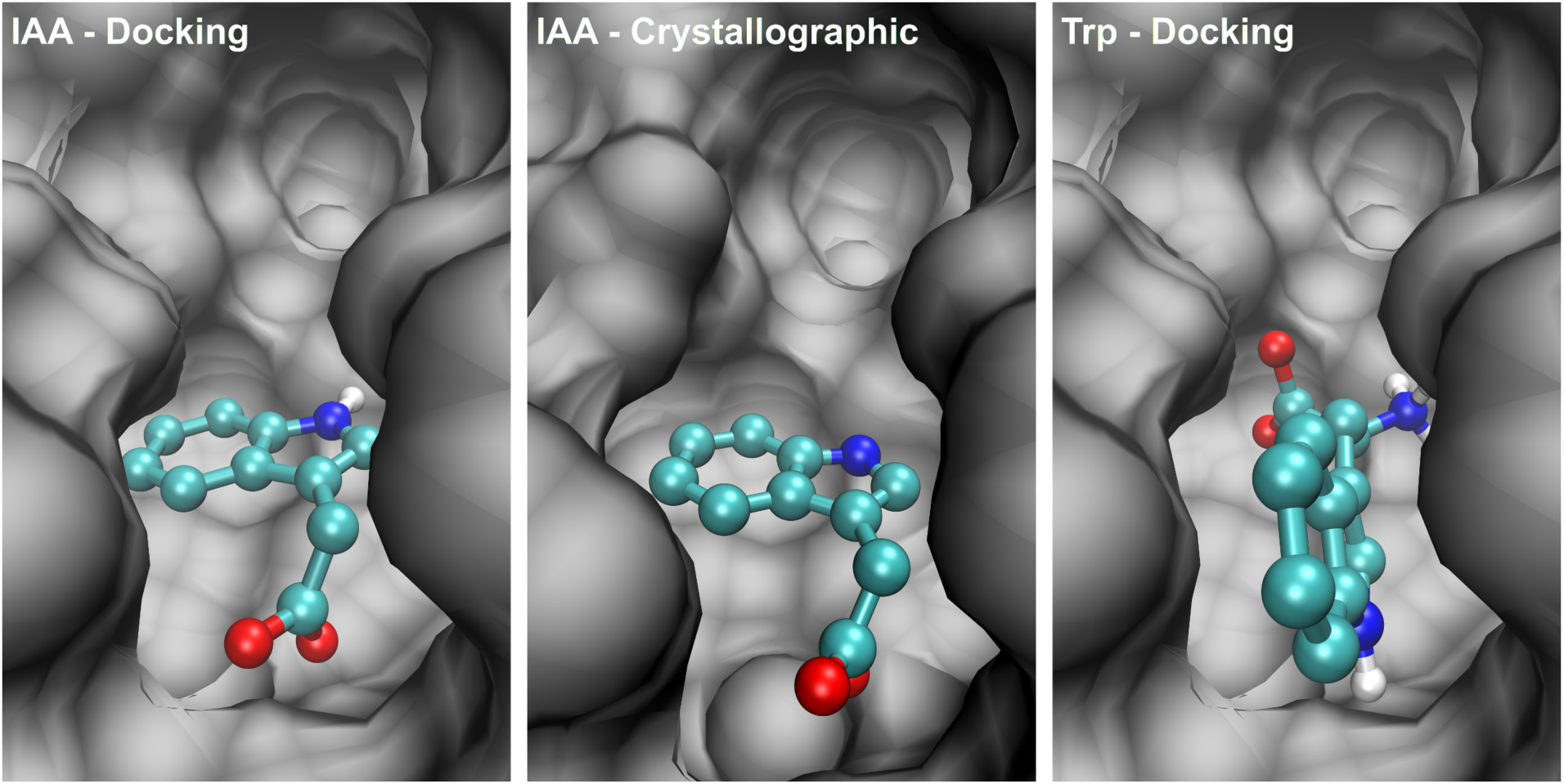
Comparison of crystallographic data and “static” docking results. The docked position of IAA at the bottom of the pocket, in the left panel, matches the crystallographic one in the centre panel. The docked position of Trp at the bottom of the pocket, in the right panel, does not match the docking and crystallographic positions of IAA.

### B. Tomographic docking

To study the transient interactions of IAA and Trp with the pocket as they move down into it, we performed tomographic docking experiments and analyzed each series of docked poses in detail, building a plausible binding pathway for both compounds over a transect of 15 Å. The docking process assigns a lower numerical score to better poses, representative of lower energy and more favourable binding. Thus, for each ligand we created a representative series of successive orientations choosing at each step the pose with the lowest score amongst all repetitions (Fig. 3). Note that the depth at which the ligand is positioned does not necessarily increase with step number. For instance, as described in greater detail below, neither the depth, nor the orientation of the docked pose of IAA changes between step 4 and step 7, indicating that the interactions relevant over these steps are dominated by the residues included at step 4, and that no further significant interactions are made until the ligand approaches residues deeper in the pocket than those at step 7. Later, we use these considerations to identify which residues are most likely to be responsible for the selection mechanism.

**Figure 3:**
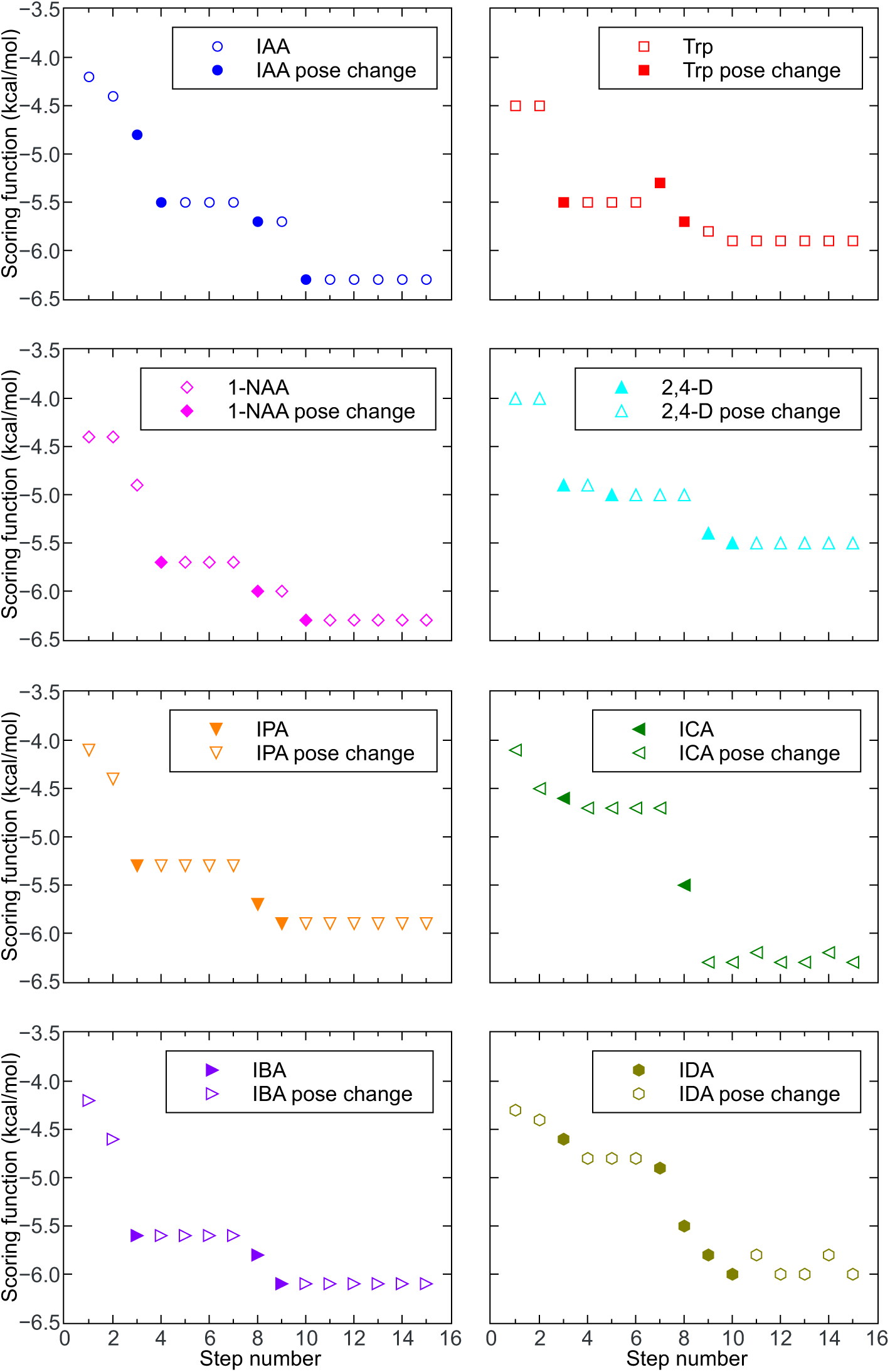
Best scores of docked poses along the transect for all compounds tested, namely IAA (blue circles), Trp (red squares) 1-NAA (magenta diamonds), 2,4-D (cyan triangles), IPA (orange triangles), ICA (green triangles), IBA (violet triangles) and IDA (olive hexagons). Each step represents the progression of the search space by 1 Å in the direction of the pocket bottom. Filled symbols indicate steps at which a significant change in depth or orientation of the docked pose occurs.

At step 1 both compounds are well out of the pocket, and at step 2 they are at the very edge of the pocket mouth. As the steps continue, the progressive inclusion of residues causes the docked position of IAA to undergo significant changes. At step 3, IAA has oriented itself with the carboxylic acid group in a niche at the back wall of the pocket (Fig. 4A). Then, for the next four steps, its scoring function, position and orientation remain constant, with the alignment of the indole perpendicular to the base of the pocket (Fig. 4B). Note that between step 3 and steps 4–7, IAA undergoes a small but significant rotation, which optimizes the perpendicularity of the indole-ring system with respect to the bottom of the pocket.

**Figure 4:**
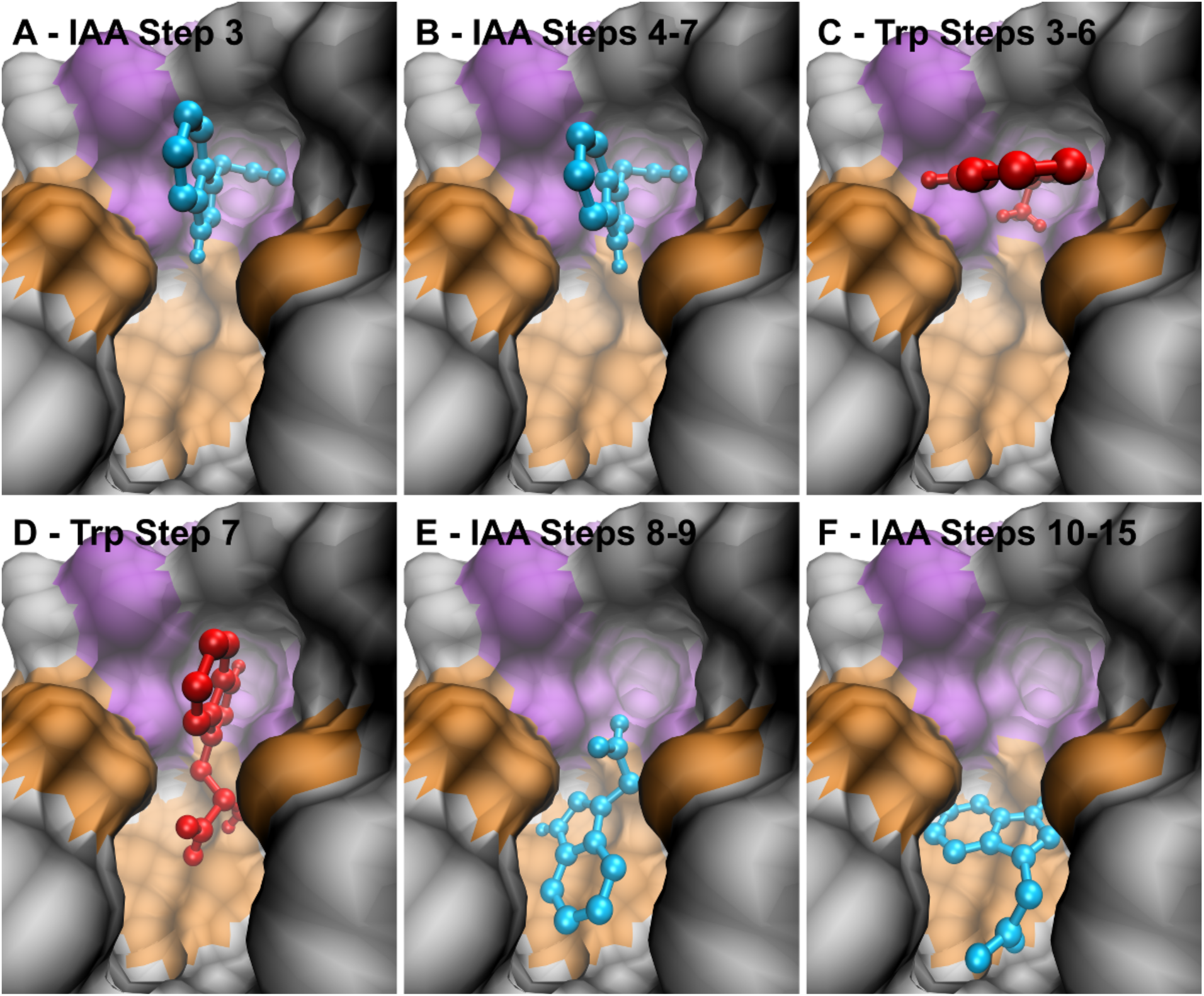
Progressive docking poses of IAA (blue) and Trp (red). The residues that form the engagement niche are highlighted in violet; those that we identify later as forming a molecular filter are highlighted in orange. (A) The position of IAA at step 3 features the side chain oriented towards the engagement niche. (B) The positions of IAA in steps 4 to 7 are superimposable, and show that the ring system is perpendicular to the bottom of the pocket. (C) The positions of Trp in steps 3 to 6 are superimposable. Its side-chain is oriented towards the niche, but the ring system is parallel, rather than perpendicular, to the pocket base. (D) At step 7, the side-chain of Trp is no longer in the niche, but points towards the pocket bottom. (E) At steps 8 and 9 (superimposable), IAA has moved towards the bottom of the pocket. (F) The positions of IAA at steps 10 to 15 are superimposable.

Considering Trp, its side-chain also becomes oriented towards the niche at step 3, and its docked depth and orientation do not change through step 6. However, unlike IAA, the orientation of the indole is parallel, not perpendicular, to the base of the pocket. This is probably due to the longer side chain and the extra rotational degree of freedom, as well as to the proximity of the aromatic rings to residues distal to the niche (Fig. 4C). The next step for Trp, step 7, presents a somersault for the pose, with its polar side groups now pointing towards the pocket bottom (Fig. 4D).

From the pose in step 7, with the tail in the niche, IAA can proceed downwards, into the position observed at step 8, via a pivoting motion of the indole from the engagement niche (cf. Fig. 4B and Fig. 4E). Poses 8 and 9 for IAA are identical, with the side-chain continuing to point towards the niche, but not in it. Then, there appears to be a final transition as residues at the base of the pocket come into play, with poses 10 to 15 showing that IAA has flipped over from poses 8 and 9 (Fig. 4F), to a position that corresponds to that found in the crystal structure [1].

### C. Binding mechanism

When the docking algorithm explores positions that include the pocket bottom, Trp is docked at the binding site. This indicates that, in principle, the final docking position is allowed. However, a detailed examination of the tomographic docking results suggests the presence of a barrier impeding the descent of the non-binder into the binding pocket, explaining why, in nature, Trp never reaches its bottom.

In the initial part of the pocket the tomographic docking identifies a region that we call the engagement niche formed by residues Lys-410, Ser-440, Gly-441, Ala-464 and Phe-465 (in violet in Fig. 5). This is the structure into which the binder orients its polar side-chain (Fig. 4B). We deem it one of the features with which potential binders need to interact in order to achieve an orientation that allows a subsequent transition to the binding position. TomoDock results suggest that the particular orientation with the ring system perpendicular to the pocket base is likely to be an essential step in the selection process. For ligands to penetrate deeper, the aromatic rings must slice down while the polar tail, anchored in the engagement niche, acts as a pivot point (cf. Fig. 4B and Fig. 4E). After this motion, the rings of IAA are positioned at the bottom of the pocket, in the vicinity of a semi-circle of non-polar residues. The ligand then undergoes a slight rotation, allowing the polar carboxylic acid group to flip and engage with the polar residues at the pocket base.

**Figure 5:**
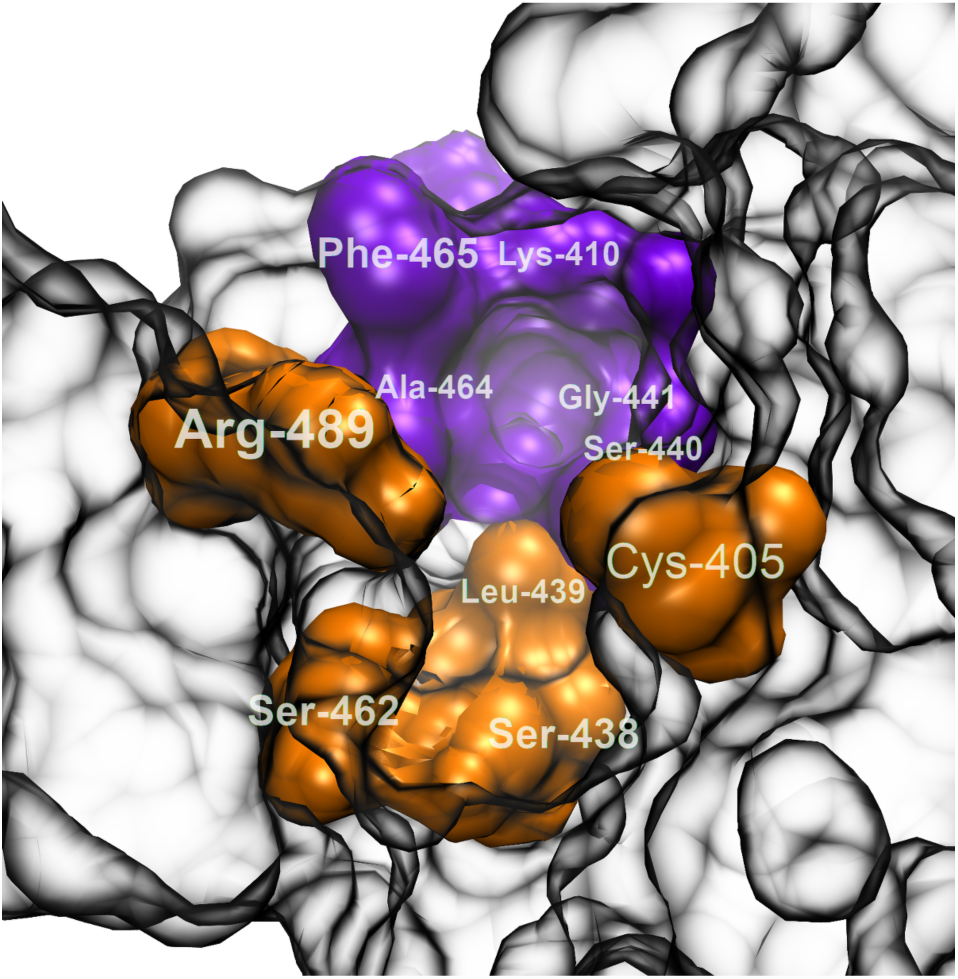
Molecular filter responsible for TIR1 selectivity. The residues shown in orange are responsible for the filtering mechanism. The engagement niche is shown in purple. The remainder of the pocket is represented with partial transparency for clarity.

To understand what blocks Trp from moving the same way as IAA, consider the results from steps 3 to 7. At step 3, Trp is docked with its polar tail in the engagement niche. However, we do not observe the perpendicular orientation of the aromatic system that we see in IAA (Fig. 4C). Note that its orientation and docking depth do not change through step 6 (Fig. 3). Then, at step 7, Trp assumes a new pose with the side-chain completely out of the niche, and pointing towards the pocket bottom (Fig. 4D). Such a geometry prevents the non-binder from moving further into the binding position via the same rotation that IAA performs, due to inappropriate orientation of the indole rings and of the side chain.

Note that the non-binder is allowed to dock further down the pocket from step 8 onwards (Fig. 3), because docking considers any position that is geometrically accessible, disregarding the motion a ligand would have to undertake in order to reach it. Also, to be active auxins, substrates need to bind in the correct orientation at the bottom of the pocket. A compound that can only interact and bind at the mouth of the pocket cannot have auxinic activity. Thus, for brevity we refer to compounds that are able to achieve an appropriate binding position simply as “binders”. Conversely, we refer to the compounds, like Trp, that cannot reach the pocket bottom as “non-binders”.

### D. Validation and experimental verification

The striking difference between IAA and Trp revealed by tomographic docking indicates that an important role is played by the residues that become available at steps 3, 4, 7, 8 and 10 (filled symbols in Fig. 3). In particular, the TomoDock results suggest that they act as a molecular filter, promoting the correct orientation of IAA, and opposing it for Trp. To identify these residues, we built a table of the atoms newly included for interaction at each step, along with the residues they belong to (Table S1). Then, we considered the new entries at the steps indicated above, taking into account the number of atoms that interact, as well as their properties.

As a first example, consider Ser-438 (see Fig. 5). This residue enters the search space at step 10, which is the first step at which IAA assumes its final binding position. Ser-438 is physically located at the bottom of the pocket and, upon close inspection, the atoms that get included at step 10 are seen to form a highly polar group. Thus, we include it in the molecular filter, and consider it responsible for the correct orientation of IAA at the pocket bottom.

As a second example, consider Ser-440. This residue is structurally part of the engagement niche, and it enters the search space at step 6, with 3 atoms. However, neither IAA nor Trp changes its position at all over this step (see Fig. 4B and C). At step 7, where 4 more atoms of Ser-440 are considered, Trp exits the niche (Fig. 4D). One could thus consider Ser-440 partly responsible for this; however, at step 7 IAA maintains the same position as it has at step 6. Given the structural similarity of the two molecules, we believe this indicates that Ser-440 does not contribute actively to ligand filtering, particularly considering that a much better candidate for the observed effect on Trp exists, namely Leu-439.

The third example we discuss is Gly-441. This is a noteworthy residue, as it is not only structurally part of the engagement niche, but also because its mutation to aspartate yields the known tir1-2 mutant [27]. This residue gives its first big conribution to the search space at step 4. But by this step both IAA and Trp have already assumed positions that do not change for a few more steps. Thus, we do not consider this residue as an active player in the molecular filter. Substitution of the large polar side group of Asp for the small non-polar Gly could interfere with binding in many ways to explain the tir1-2 phenotype.

Performing this analysis on all viable residues shows that the filter is formed by Cys-405, Ser-438, Leu-439, Ser-462 and Arg-489 (in orange in Fig. 5). Of these, Arg-489 seems to be the residue that most significantly affects the orientation of the compounds with respect to the engagement niche, of which, however, it is not a part. Leu-439 and Ser-462 appear to block the descent of Trp and promote that of IAA, as they progressively enter the search space in steps 7–9. Finally, Cys-405 and Ser-438 are likely to be instrumental for IAA to assume the final binding position, since they start contributing significantly at steps 8 and 10, respectively. Note that all the residues in physical proximity of the ligand at the pocket bottom are likely to be involved in stabilizing docked auxins, but they are not necessarily part of the filtering mechanism.

To further exemplify our method, we apply it to six other potential binders (Fig. S2), namely indole-3-carboxylic acid (ICA), indole-3-propionic acid (IPA), indole-3-butyric acid (IBA), indol-3-yl acetate (IDA), 1-naphthaleneacetic acid (1-NAA) and 2,4dichlorophenoxyacetic acid (2,4-D). The aliphatic side-chains of ICA, IPA and IBA differ in length from that of IAA, being one atom shorter, one atom longer and two atoms longer, respectively. Esterifying indole-3-ol with acetic acid gives IDA, while changing the indole system to a naphtalene double ring yields 1-NAA. Finally, we include 2,4-D as it is one of the most widely used herbicides in the world. For all compounds, tomographic docking predicts a plateaux between steps 3 and 7 (Fig. 3). At these steps, all compounds are docked at the right depth to permit interaction with the engagement niche. Results for 1-NAA, 2,4-D and IPA show that they do engage with the niche, with the aromatic ring system in the same position as that of IAA, perpendicular to the base of the pocket (Fig. 6A–C). In contrast, the results for ICA, IBA and IDA predict that they are not able to adopt the correct engagement pose to allow the subsequent pivot. Whilst they may occupy space in and interact with the outer chamber, TomoDock suggests that they fail to orient appropriately for transit further down the pocket. In the case of ICA and IDA, there is no interaction with the engagement niche, while IBA orients its ring systems transversal, rather than perpendicular, to the pocket bottom, similar to Trp (Fig. 6D–F). These poses are consistent with 1-NAA, 2,4-D and IPA being active ligands for TIR1, albeit with different affinities, and with ICA, IBA and IDA being not.

**Figure 6:**
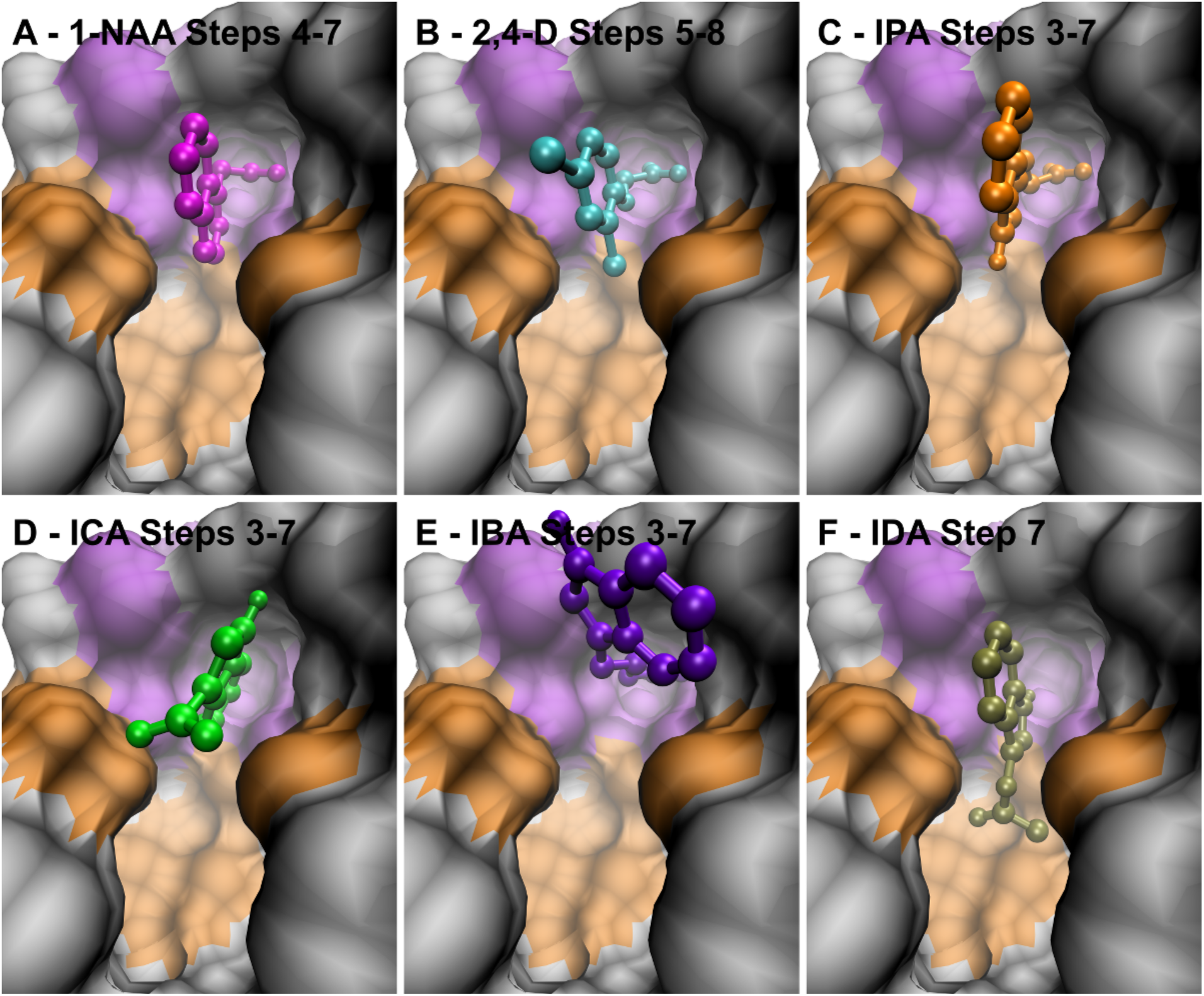
Docking poses of 1-NAA (magenta), 2,4-D (cyan), IPA (orange), ICA (green), IBA (violet) and IDA (olive). (A) The positions of 1-NAA at steps 4 to 7 (superimposable) show the side-chain of the ligand engaged with the niche, and its ring system perpendicular to the pocket bottom. (B) In steps 5 to 8 (superimposable) the orientation of 2,4-D is the same as that of IAA and 1-NAA. (C) At steps 3 to 7, also IPA engages the niche with the ring system perpendicular to the pocket base. The positions at these steps are superimposable. (D) In steps 3 to 7 (superimposable), the tail of ICA never engages the niche. (E) The tail of IBA finds the niche in steps 3 to 7 (superimposable), but the ring system is misoriented. (F) At step 7, the side-chain of IDA is oriented towards the pocket bottom, in a pose reminiscent of that of Trp at the same depth.

Experimental confirmation of binding from SPR and root growth assays, (Fig. 7 and Tab. I), support the numerical predictions. In particular, SPR experiments indicate that Trp, ICA, IBA and IDA have no measurable binding activity at 50 *µ*M, a concentration far in excess of the IC50 value of all active auxins (Tab. I). IPA and 2,4D, instead, bind weakly compared to IAA and 1-NAA, with noticeably more rapid dissociation rates (see Fig. 7 and related results in [2]). Like IAA, 1-NAA is a strong ligand. Notice that, as mentioned above, substrate binding to the receptor is only the first step in the auxinic interaction, and the binding of the co-receptor IAA7 (in our case) to the substrate-receptor complex can only happen if the substrate is bound in the bottom of the receptor pocket, and in the correct orientation. Thus, SPR experiments offer a good method to validate the computational results: if a substrate binds incorrectly, it will impair or entirely preclude the binding of the receptor to the IAA7 co-receptor and produce no SPR signal. The relative effectiveness of each compound in root growth assays compares favourably with the SPR measurements. Trp, ICA, IBA and IDA inhibit root growth only at very high concentrations, where phytotoxicity sets in. IPA is seen to be a weak auxin. The root growth assays suggest that 2,4-D is the most active auxin, more active than the SPR data suggest. However, root growth assay activities depend on tissue and cellular transport, as well as on receptor binding of the compounds in question, and so IC50 values do not correspond exactly to in vitro binding values. Nonetheless, the assays are still a useful verification method, as one cannot observe root-growth inhibition for compounds that do not correctly bind to the receptor.

**Figure 7:**
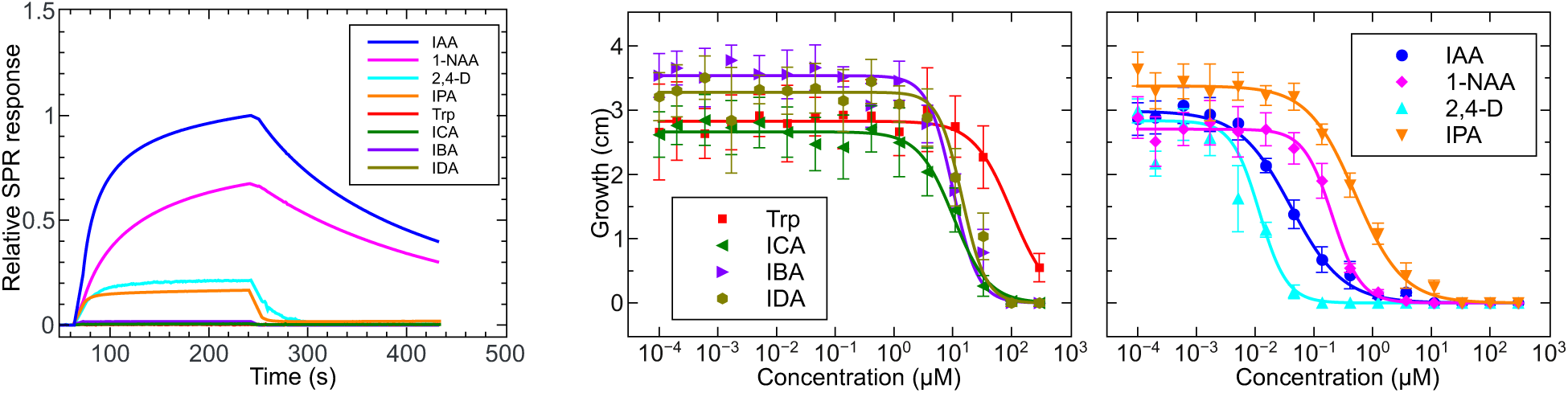
Surface plasmon resonance and root growth assay results confirm the numerical predictions. The results from SPR experiments (left panel) show that Trp (red line), ICA (green line), IBA (violet line) and IDA (olive line) do not bind TIR1 at all. Conversely, 1-NAA (magenta line), 2,4-D (cyan line) and IPA (orange line) all bind, although with differing activities from IAA (blue line). All compounds were tested at 50 *µ*M. Root growth inhibition measurements (centre and right panels) substantially confirm these results, revealing that Trp (red squares), ICA (green triangles), IBA (violet triangles) and IDA (olive hexagons) do not inhibit root growth up to extreme concentrations. In the right-hand panel, 1-NAA (magenta diamonds), 2,4-D (cyan triangles) and IPA (orange triangles) are all active auxins. Derived IC50 values are given in Table I.

**Table I:** IC50 for primary root growth inhibition derived from root growth assays.

| Compound | $\text{IC}_{50}$ ($\mu\text{M}$) |
| --- | --- |
| 2,4-D | $0.0118 \pm 0.0009$ |
| IAA | $0.041 \pm 0.007$ |
| 1-NAA | $0.206 \pm 0.015$ |
| IPA | $0.51 \pm 0.06$ |
| IBA | $10.1 \pm 1.2$ |
| ICA | $10.4 \pm 1.2$ |
| IDA | $15 \pm 2$ |
| Trp | $102 \pm 13$ |

## IV. CONCLUSIONS

We have studied the binding process of the plant growth hormone IAA onto its main receptor TIR1, to investigate its selectivity mechanism. For our study, we developed a novel tomographic docking approach suitable for investigating deep binding pockets in a series of pseudo-time steps. The method gradually changes the search space of a docking algorithm to allow one to consider sequential interactions of each potential ligand with pocket residues at increasing depths. This mimics what happens in nature when a small molecule descends into a binding cavity. In the present study we have considered the receptor structure as rigid, as is the case for most docking experiments. However tomographic docking can be adapted to allow for the flexibility of some side-chains of the receptor, and this will be the subject of future work. The tomographic method shows a plateau of scoring function values part-way down the pocket, indicating a region over which transient interactions are made en route to the docking site at the base.

Detailed study of the docking poses obtained for the natural ligand IAA and the related, but non-auxin Trp over this region points towards two features in the pocket responsible for selectivity. The first, an engagement niche in the back of the pocket, allows potential ligands to orient before subsequent motion towards the binding site. The second, a molecular filter, promotes the correct pose of the aromatic ring system for binders, necessary to access the pocket bottom. Tryptophan and a set of non-binders assume sub-optimal orientations, and are prohibited from onwards motion.

The identification of the residues that form the engagement niche and the molecular filter makes them a fundamental study subject for the rational design of novel auxin-based herbicides. They are critical for selectivity, and constitute preferred mutation targets for further experiments. One such mutant, tir1-2 (G411-Asp), is already known, and experiments have shown that the mutation has a small but measurable effect on the resistance of the plant to the auxin-transport inhibitor 2-carboxyphenyl-3-phenylpropane-1,2-dione (CPD) [27], although this non-conservative substitution may not help elucidate the role of residue 441 further.

Experimental results from SPR and root growth assays performed on a set of active and inactive compounds are consistent with TomoDock results, confirming the validity of our method. The application of tomographic docking need not be limited to the analysis of auxin binding to TIR1. In fact, it can be used to examine also other members of the TIR/AFB auxin receptor family. Identifying similarities and differences between the interaction mechanisms in different receptors can play a key role in designing receptor-specific compounds, which are very useful in controlling herbicide resistance. In addition, the tomographic docking principle is general, and can be applied to any deep binding site. Thus, proteins such as those involved in the transport of small molecules, as well as enzymes and channel proteins, are all natural targets for tomographic docking investigation.

### Data accessibility

The protein structures used in this study are available from the Protein Data Bank web site at the URL http://www.rcsb.org/pdb/home/home.do The TomoDock code is available on the web pages of the authors.

### Authors’ contributions

VVU and CIDG performed the numerical simulations. MQ designed and performed all the SPR and bioassay experiments. CIDG implemented the TomoDock code. All authors developed the principle of tomographic docking, from an original idea of VVU. All authors analyzed data and results, and wrote the manuscript.

### Funding

VVU and RMN acknowledge support from BBSRC via grant n. BB/L009366. MQ acknowledges support from BBSRC via a studentship awarded through the Midlands Integrative Biosciences Training Partnership (MIBTP).

## References

[1] Tan X, Calderón-Villalobos LIA, Sharon M, Zheng CX, Robinson CV, Estelle M, Zheng N. 2007 Mechanism of auxin perception by the TIR1 ubiquitin ligase. Nature 446 640–645. (doi:10.1038/nature05731)

[2] Calderón Villalobos LIA et al. 2012 A combinatorial TIR1/AFB-Aux/IAA co-receptor system for differential sensing of auxin. Nat. Chem. Biol. 8, 477–485. (doi:10.1038/nchembio.926)

[3] Napier RM. 2014 Auxin Receptors and Perception. In Auxin and Its Role in Plant Development. E. Zażímalová, J. Petrášek and E. Benková, editors. Springer-Verlag, Vienna, 101–116. (doi:10.1007/978-3-7091-1526-8)

[4] Napier RM. 2003 Regulators of growth: Auxins. In Encyclopedia of Applied Plant Sciences. B. Thomas, D. J. Murphy, B. G. Murray, editors. Academic Press, Waltham, MA, 985–995.

[5] Lee S, Sundaram S, Armitage L, Evans JP, Hawkes T, Kepinski S, Ferro N, Napier RM. 2014 Defining Binding Efficiency and Specificity of Auxins for SCF^TIR1/AFB^- Aux/IAA Co-receptor Complex Formation. ACS Chem. Biol. 9 673–682. (doi:10.1021/cb400618m)

[6] Kitchen DB, Decornez H, Furr JR, Bajorath J. 2004 Docking and scoring in virtual screening for drug discovery: Methods and applications. Nat. Rev. Drug Discov. 3 935–949. (doi:10.1038/nrd1549)

[7] Jorgensen WL 2004 The many roles of computation in drug discovery. Science 303 1813–1818. (doi:10.1126/science.1096361)

[8] Wishart DS, Knox C, Guo AC, Cheng D, Shrivastava S, Tzur D, Gautam B, Hassanali M. 2008 DrugBank: a knowledgebase for drugs, drug actions and drug targets. Nucleic Acids Res. 36 D901–D906. (doi:10.1093/nar/gkm958)

[9] Stark JL, Powers R. 2012 Application of NMR and Molecular Docking in Structure-Based Drug Discovery. In NMR of Proteins and Small Biomolecules. G. Zhu, editor. Springer-Verlag, Berlin, 1–34. (doi:10.1007/128_2011_213)

[10] Xie Z-R, Hwang M-J. 2015 Methods for Predicting Protein-Ligand Binding Sites. In Molecular Modeling of Proteins. A. Kukol, editor. Springer Science, New York, NY, 383–398. (doi:10.1007/978-1-4939-1465-4_17)

[11] Kobayashi S, Boggon TJ, Dayaram T, Janne PA, Kocher O, Meyerson M, Johnson BE, Eck MJ, Tenen DG, Hal-mos B. 2005 EGFR mutation and resistance of non-smallcell lung cancer to gefitinib. New Engl. J. Med. 352 786–792. (doi:10.1056/NEJMoa044238)

[12] Bologa CG et al. 2006 Virtual and biomolecular screening converge on a selective agonist for GPR30. Nat. Chem. Biol. 2 207–212. (doi:10.1038/nchembio775)

[13] Choi JS, Braymer JJ, Nanga RPR, Ramamoorthy A, Lim MH. 2010 Design of small molecules that target metal-A^ species and regulate metal-induced. Aft aggregation and neurotoxicity. Proc. Natl. Acad. Sci. USA 107 21990–21995. (doi:0.1073/pnas.1006091107)

[14] Smith CC et al. 2012 Validation of ITD mutations in FLT3 as a therapeutic target in human acute myeloid leukaemia. Nature 485 260–U153. (doi:10.1038/nature11016)

[15] Hyster TK, Knorr L, Ward TR, Rovis T. 2012 Biotinylated Rh(III) Complexes in Engineered Streptavidin for Accelerated Asymmetric C-H Activation. Science 338 500–503. (doi:10.1126/science.1226132)

[16] Xu W, Lucke AJ, Fairie DJ. 2015 Comparing sixteen scoring functions for predicting biological activities of ligands for protein targets. J. Mol. Graphics and Model. 57 76–88. (doi:10.1016/j.jmgm.2015.01.009)

[17] Trott O, Olson AJ. 2010 AutoDock Vina: improving the speed and accuracy of docking with a new scoring function, efficient optimization and multithreading. J. Comput. Chem. 31 455–461. (doi:10.1002/jcc.21334)

[18] AutoDock Vina http://vina.scripps.edu/

[19] Schindelin J, Rueden CT, Hiner MC, Eliceiri KW. 2015 The ImageJ ecosystem: An open platform for biomedical image analysis. Mol. Reprod. Dev. 82 518–529.

[20] ImageJ http://imagej.net/

[21] Humphrey W, Dalke A, Schulten K. 1996 VMD - Visual Molecular Dynamics. J. Molec. Graphics. 14 33–38. (doi:10.1016/0263-7855(96)00018-5)

[22] VMD - Visual Molecular Dynamics http://www.ks.uiuc.edu/Research/vmd/

[23] Morris GM, Huey R, Lindstrom E, Sanner MF, Belew RK, Goodsell DS, Olson AJ. 2009 Autodock4 and AutoDockTools4: automated docking with selective receptor flexibility. J. Comput. Chem. 16 2785–2791. (doi:10.1002/jcc.21256)

[24] AutoDock http://autodock.scripps.edu/resources/adt

[25] Woodward AW, Bartel B. 2005 Auxin: Regulation, action, and interaction. Ann. Bot. - London 95 707–735. (doi:10.1093/aob/mci083)

[26] Lin CY, George Mw, Gill PMW. 2004 EDF2: A Density Functional for Predicting Molecular Vibrational Frequencies. Aust. J. Chem. 57 365–370. (doi:10.1071/CH03263)

[27] Ruegger M, Dewey E, Gray WM, Hobbie L, Turner J, Estelle M. 1998 The TIR1 protein of Arabidopsis functions in auxin response and is related to human SKP2 and yeast Grr1p. Gene. Dev. 12 198–207. (doi:10.1101/gad.12.2.198)

